# Third-generation sequencing and the future of genomics

**DOI:** 10.1101/048603

**Authors:** Hayan Lee, James Gurtowski, Shinjae Yoo, Maria Nattestad, Shoshana Marcus, Sara Goodwin, W. Richard McCombie, Michael C. Schatz

## Abstract

Third-generation long-range DNA sequencing and mapping technologies are creating a renaissance in high-quality genome sequencing. Unlike second-generation sequencing, which produces short reads a few hundred base-pairs long, third-generation single-molecule technologies generate over 10,000 bp reads or map over 100,000 bp molecules. We analyze how increased read lengths can be used to address longstanding problems in *de novo* genome assembly, structural variation analysis and haplotype phasing.

## Introduction

Since the advent of “second-generation” sequencing (or next-generation sequencing) with the commercialization of Roche/454 pyrosequencing in 2005, Illumina/Solexa sequencing in 2007, and other high-throughput technologies, the cost of genome sequencing has precipitately dropped^1^. This has enabled the sequencing of many new genomes^2^ along with widespread resequencing efforts to analyze genomic diversity^3^. Although second-generation sequencing has enabled population-scale analyses of single nucleotide and other small variants, analysis of larger structural variations has proved difficult. Further, new genomes assembled *de novo* using second-generation technologies are often of lower quality compared with those genomes sequenced using older, more expensive methods^4, 5^. In particular, *de novo* genome assemblies comprised only of short-reads can lack entire portions of genomes, may be fragmented and missing important genes, and lack sufficient robustness to study overall chromosome architecture^2, 6^. In some cases the assembled sequences have been substantially smaller than the average gene size rendering the sequence less useful than earlier reference genomes^7^. Resequencing projects have also been severely limited in their analysis of structural variations, missing tens of thousands of structural variants or more per mammalian-sized genome^8^.

The availability of new single-molecule sequencing technologies that can produce average read lengths of more than 10,000bp and some read lengths up to 100,000bp or more **(Table 1)** has enabled greatly improved analysis of genome structure. Importantly, longer read lengths span more repetitive elements and thus produce more contiguous reconstructions of the genome^9^. With respect to structural variation analysis, long reads enable improved “split-read” analyses so that insertions, deletions, translocations and other structural changes can be more readily recognized^8^. Furthermore, the singlemolecule sequencing technologies often produce more uniform coverage of the genome, as they are not as sensitive to GC content as second generation technologies which tend to have reduced or completely absent coverage over regions with imbalanced sequence composition^10^. Complementary to the improved sequencing technologies, several long-range mapping technologies are available that can map the structure 50kbp to 250kbp or longer molecules using florescent probes and other markers. Using third-generation sequencing and mapping technologies together it is possible to form super-contigs (“scaffolds”) that can span nearly entire chromosome arms leading to greatly improved structural analysis^11–13^.

**Table 1:** Characteristics of 3^rd^ generation DNA sequencing and mapping platforms. Contig/Scaffold N50 indicates the N50 length of the de novo assembled contigs/scaffolds. Haplotype phasing indicates the N50 length of the phased regions of the genome. N50 size is a weighted median average: half of the total sequence length has been resolved into sequences this size or longer. ^*^Includes the cost and time for both sample preparation and short read sequencing using a NextSeq/HiSeq2500. †Assumes library construction and instrument costs can be amortized over multiple runs. All prices subject to change, see https://www.dugsim.net/estimate_cost for current estimates.

|  | ***Illumina TruSeq***<br>***Synthetic Long Reads*** | ***Pacific***<br>***Biosciences*** | ***Oxford Nanopore***<br>***Technologies*** |
| --- | --- | --- | --- |
| Technology | Barcoded & Amplified<br>Synthetic long reads | Single Molecule<br>Real Time Sequencing | Nanopore<br>Sequencing |
| Mean Length | 3-5kbp | 10-15kbp | 5-10kbp |
| Raw Error Rate | 0.1% | 10-15% | 10-30% |
| Costs / GB | ~\$2500* | ~\$500† | ~\$1000† |
| Time / GB | 2-3 days* | 2-3 hours | 1-2 days |
| Human Metrics | 0.5Mbp<br>Haplotype phasing N50<sup>19</sup> | 26.9 Mbp<br>Contig N50<sup>46</sup> | N/A |

**Table 1: Characteristics of 3<sup>rd</sup> generation DNA sequencing and mapping platforms.**
|  | ***BioNanoGenomics*** | ***10X Genomics*** | ***Dovetail cHiCago*** |
| --- | --- | --- | --- |
| Technology | Optical mapping of<br>fluorescent probes | Barcoded<br>“Read Clouds” | Chromatin<br>mate-pairs |
| Mean Span | 100-250kbp | 30-100kbp | 25-100kbp |
| Error Modes | Fragile sites,<br>incomplete labeling | Barcode reuse,<br>Short read mapping | Variable span,<br>short read mapping |
| Costs / Mammalian | ~\$3,000 | ~\$2,000* | ~\$10,000* |
| Time / Mammalian | 1-2 days | 2-3 days* | 2-3 weeks* |
| Human Metrics | 31.1 Mbp<br>Scaffold N50<sup>20</sup> | 21.6 Mbp<br>Haplotype phasing N50<sup>47</sup> | 29.9 Mbp<br>Scaffold N50<sup>30</sup> |

Third-generation technologies have been used to produce highly accurate *de novo* assemblies of hundreds of microbial genomes^14, 15^ and highly contiguous reconstructions of many dozens of plant and animal genomes, enabling new insights into evolution and sequence diversity^16–18^. They have also been applied to resequencing analyses, to create detailed maps of structural variations^8^ and phasing variants^19^ across large regions of human chromosomes. Notably, the new technologies have been used to fill in many of the gaps in the human reference genome that had resisted more than one decade of scrutiny^8, 20^. One clinically important application of the improved read lengths is to sequence medically relevant regions of the genome, such as the human leukocyte antigen (HLA) genes of the major histocompatibility complex^21^. The technologies have proven instrumental for resolving the composition of metagenomics communities, as longer read lengths and longer spans allow for the assembly of individual species within mixtures too complex to be resolved by short reads alone^22^. Outside of DNA sequencing, third-generation technologies have also been widely used to study transcriptomes, recognizing thousands of novel isoforms and gene fusions that were not found using second-generation short read sequencing^23^. Finally, some of the technologies also allow for direct measurement of epigenetic modifications from single molecules, allowing for many new methyltransferases to be discovered and for the role of methylation in pathogens to be better studied^24^.

Here we analyze the capabilities of third-generation technologies to show how they improve the “3Cs of Genome Sequencing”: the contiguity, completeness and correctness of a genome. We discuss the key characteristics of the technologies and the analysis algorithms needed to effectively use them. We then undertake a meta-analysis of the currently available 3^rd^ generation genome assemblies, a retrospective analysis of the development of the reference human genome, and simulations with dozens of species across the tree of life. From these data, we develop a new predictive model of genome assembly presented as an online web-service (http://qb.cshl.edu/asmmodel/predict.html) that can accurately estimate the performance of a genome assembly project using different technologies **(Online Methods)**.

## Third-generation sequencing

The three commercially available third-generation DNA sequencing technologies are Pacific Biosciences (PacBio) Single Molecule Real Time (SMRT) sequencing, the Illumina Tru-seq Synthetic Long-Read technology and the Oxford Nanopore Technologies sequencing platform. Using single-molecule sequencing or clonal amplification and sequencing of long molecules, all three technologies can produce long reads averaging between 5,000bp to 15,000bp, with some reads exceeding 100,000bp.

The most established of these is the PacBio SMRT technology, which was commercially introduced in 2010^9^. The SMRT technology sequences DNA using sequencing-by-synthesis, and optically monitors fluorescently tagged nucleotides as they are incorporated into individual template molecules. The current instrument, the PacBio RS II, produces read lengths of up to ~100,000 bp with the greatest throughput (~8GB / day) of the currently available long-read technologies **<u>(Table 1)</u>**. Their recently released PacBio Sequel instrument is poised to increase the throughput by as much as 7 fold. Reads have a raw error rate of 10% to 15% but nevertheless several algorithmic techniques have been developed that can improve the per-nucleotide accuracy to over 99.99% or more with sufficient coverage (Online Methods Table 1). Approximately 50x long read coverage is required for “self-correction” approaches, while lower coverage can be effectively used with hybrid error correction algorithms that leverage additional high coverage short read sequencing to error correct the long reads^16^. The main limitation with PacBio sequencing is the cost relative to second-generation approaches, which has limited its application for analyzing large numbers of genomes. Nevertheless, to date hundreds of projects have successfully used PacBio sequencing, including nearly perfect assemblies or very high quality genomes of microbes, fungi, plant and animal species, as well as very high quality *de novo* assemblies of entire human genomes^16^.

The second third-generation technology, introduced in 2012, was the Moleculo protocol which is now marketed as Illumina TruSeq Synthetic Long Reads^19^. Using this approach, ~10kbp molecules of DNA are clonally amplified and barcoded before sequencing with a short read instrument, so that long reads can be synthetically created from the short read sequences. The synthetic long reads are very accurate (~0.1% error) **(Table 1)**, and can be used for phasing analyses and assembly without error correction. However, because TruSeq relies on long-range amplification and the reads are synthetically generated, the available read lengths are shorter than other approaches, and are prone to termination and biases in any region where the Illumina chemistry is biased, such as regions with high GC content or tandem repeats. Finally, obtaining sufficient coverage for *de novo* genome assembly can be expensive, often even greater than PacBio sequencing, since 900x to 1500x or more short read coverage may be required to assemble 30x coverage of synthetic long reads. Nevertheless, several studies have used the technology for assembling and phasing complex genomes, including phasing very large regions of human chromosomes^19^.

The most recent third-generation technology was released by Oxford Nanopore Technologies in 2014. Their current instrument, the Oxford Nanopore MinION is a handheld device that sequences DNA by electronically measuring the minute disruptions to electric current as DNA molecules pass through a nanopore^15^. The read lengths of the currently available instrument are similar to those produced by PacBio **(Table 1)**, although to date the instrument has suffered from worse accuracy and lower throughput which has limited it’s scope to sequencing small genomes, including E. coli (4.5Mbp) or yeast (12Mbp), or amplicons. Using error correction algorithms similar to those that are available for PacBio reads, the per-nucleotide accuracy of genomes sequenced using the MinION has been measured to be >99.95% ^15^. Interestingly, the instrument’s small size and low cost have empowered it to be used for studies in very remote locations, including studying Ebola outbreaks in the field in West Africa^25^.

## Third generation mapping

Mapping technologies determine the large-scale sequence structure of DNA without sequencing every base. One of the original mapping technologies, genetic maps, are constructed by analyzing the recombination rates between heterozygous markers. This requires genotyping large populations, which may not be available for some species and is labor intensive to collect^26^. More recent second-generation mapping technologies include “mate-pair” libraries, which comprise pairs of reads separated by a known span with unsequenced bases in between^27^. The mate-pair approach is commonly used to produce “jumping” libraries with pairs of reads separated by a few kilobases, but is less reliable for mapping loci that are further apart unless additional constructs such as fosmids or BACs are used^28^. For example, a mate-pair approach was used to assemble a diploid human genome *de novo* with a contig N50 size of 484kbp^28^, far better than is possible using short-reads alone, which typically produce human assemblies with contig N50 sizes of at most 20-30kbp. However, >20,000 fosmid pools were required, and each had to be individually prepared and sequenced, meaning that whilst accurate, significant labor and sequencing costs were incurred that limit widespread adoption of this method.

One of the most successful third-generation mapping systems available is the Irys system from BioNano Genomics, launched in 2010. The Iris is an optical mapping system using fluorescently tagged probes attached at “nicked” restriction digest sites to fingerprint long DNA molecules **(Table 1)**^11^. After imaging, the per-molecule fingerprints are assembled into larger optical maps, typically spanning many megabases of a chromosome. Irys maps can be compared to a sequence assembly to construct scaffolds of how the sequences should be ordered and oriented along the chromosome, or compared to a reference genome on their own to reveal structural changes, such as the rearrangement or fusion of two chromosomes. Irys suffers from biases that have limited its use, especially incomplete nicking of the DNA, causing a proportion of the digest sites remain unlabeled, and “fragile sites” where multiple nick sites in close proximity cause the DNA to systemically shear and limit the overall length of the map. Nevertheless, several studies have used Irys data together with second or third-generation sequencing technologies to improve scaffolding and structural resolution^13, 29^. Notably, a combination of PacBio reads and Irys mapping produced one of the most contiguous de novo assemblies of a human genome to date, with a 1.4Mbp contig N50 and a 31.1 scaffold N50^13^ reveal many hundreds of novel structural variations.

Other third-generation mapping protocols have been developed to construct long-range mate-pair-like reads from chromatin interactions. In the early studies, chromatin interactions measured via the “Hi-C” protocol were used as very long range, if variable length, mate-pairs that spanned hundreds of kilobases or more^12^. Because most chromatin interactions are highly localized, the relative order and orientation of assembled contigs can be inferred from the density of Hi-C mappings. More recently, Dovetail Genomics introduced an optimized Hi-C approach in spring 2015 called the cHiCago protocol, which crosslinks DNA within artificial constructs that limit transient long-range or inter-chromosomal interactions^30^. Mate-pair like cHiCago data can map long spans using relatively inexpensive reagents and second-generation reads. This technology is proprietary to Dovetail, and samples must be shipped and processed on site, which could limit their potential application.

The newest third-generation mapping technology is the Chromium instrument from 10X Genomics, introduced spring 2016. It is conceptually similar to the Illumina TruSeq Synthetic Long Read approach, but uses oil emulsion and multiple displacement amplification (MDA) to amplify and ligate short barcode sequences across much longer molecules (>100kbp). However, because the short reads are sequenced to very low coverage (~0.1x per molecule) they cannot be assembled into ‘synthetic’ long reads. Instead each barcode demarcates a “read cloud” of short reads that are localized in the genome. The read cloud information is used to scaffold *de novo* assemblies, structural variation analysis, and haplotype phasing, including phasing megabase regions of the human genome^31^.

#### BOX: Third-generation genomics algorithms

**Genome assembly** The fundamental obstacle to a high quality genome assembly is repetitive sequences. Whilst second-generation short sequencing reads can be used to assemble long non-repetitive sequences, short-read assemblers are fundamentally unable to assemble repeat sequences that are longer than the available read (or span) length^16^. Third-generation long sequencing reads have proved invaluable for achieving high quality assemblies because they span proportionally more of the repeats present in a genome.

Long read assemblers use an *overlap graph* or *string graph* approach that begins by comparing the long reads in their entirety to each other. Owing to their high quality, raw Tru-seq reads can be used directly by assembly algorithms. However, because of the high frequency of errors, both PacBio and MinION sequencing reads must be pre-processed either with *hybrid error correction*, which uses an alignment of high quality short-read data to error-correct the long reads^32^, or *self-correction,* in which the long reads are aligned to each other to form an error corrected consensus sequence^16, 33^. Hybrid strategies are more effective when a limited amount of long read coverage is available, especially below 30x coverage, whereas self-correction is better suited to higher sequencing coverage because more reliable alignments can be made between the long reads. Third-generation sequencing technologies produce a “log-normal” read length distribution, which introduces a long tail of read lengths to include some over 100kbp. This distribution also implies deep coverage (50x to 100x or more) is needed for the best possible assembly, since a large fraction of the data will consistent of relatively short reads (<1kbp). Furthermore, deep coverage increases the availability of the longest possible reads. For example, with only 30x coverage of current PacBio long reads averaging 10kbp, approximately 5x coverage of reads longer than 20kbp will be available, but doubling the overall coverage will lead to 10x coverage of reads more than 20kbp. Those ultra-long reads are the most useful for resolving repeats, and their use improves contig sizes and quality.

Algorithm development for third-generation technology continues. For example, the recently published MHAP self-correction algorithm uses a clever hashing strategy to align PacBio reads to each other and produce an error corrected version of each read^16^. Then the error corrected PacBio reads can be assembled using a long read genome assembler, such as the Celera Assembler that uses a string-graph approach. Using MHAP, Koren *et al*. assembled 5 genomes to very high quality, including *E. coli*, yeast *S. cerevisiae*, *D. melanogaster*, *A. thaliana*, and the CHM1 hydatidiform mole human genome^16^. In the case of Oxford Nanopore Technologies, the published reports have included near perfect microbial and yeast genomes sequenced using various self-correction or hybrid approaches^15^. For Illumina TruSeq sequencing, with read lengths averaging approximately half that of PacBio or Oxford Nanopore, and consequently the best *de novo* assemblies of large eukaryotic genomes have achieved contig N50 sizes of a few hundred kilobases^34^.

**Chromosome scaffolding** Third-generation mapping technologies order and orient contigs into larger scaffolds, using either ‘greedy’ approaches that iteratively link together contigs with the strongest linking support, or by a ‘global optimization’ that tries to best satisfy all of the linking information at once^30^. Combining sequencing with long-range mapping data cannot only improve assemblies but is also potentially more cost effective. For example, the Dovetail cHiCago approach combines their mapping technology with a *de novo* assembly of relatively inexpensive Illumina short-reads.

One of the biggest challenges for chromosome scaffolding is obtaining a sufficiently high quality sequence assembly before scaffolding: for BioNano Genomics, this is required to have several nick sites on each contig so that the optical map can be confidently aligned; and for 10X and Dovetail this is needed to detangle the initial read cloud or chromatin mate-pair information. In particular the Dovetail human scaffolding result began with a 178kb short-read assembly scaffold N50 size, the best BioNano Genomics human results began with a 1.4Mbp contig N50 size, and Hi-C approaches are very limited if the scaffold N50 length is below 50kbp^12^. The success of these technologies is also very sensitive to any biases in the data; BioNano Genomics map data are limited by fragile sites, and the Dovetail cHiCago protocol was designed to filter out the biological noise of chromatin domains from the desired technical signal of locality. Dovetail cHiCago and 10X genomics will also be biased by the limitations of Illumina sequencing, especially reduced coverage in regions with extreme GC content. Finally and most significantly, scaffolding a chromosome has less information than fully sequencing a chromosome, and important biological sequences could be missed or obscured in the gaps between the contigs.

**Structural variation analysis.** Finding SNPs and small variants is now relatively straightforward with short reads, but detecting structural variations (variations >50bp) is more difficult because short reads tend to fail to map to the breakpoints of a structural variant^35^. Third-generation mapping and sequencing enable improved *split-read* analysis since splitting a 10kbp long read or 100kb optical map in half still allows for 5kbp or 50kbp to be confidently aligned^8^. Many structural variations are also flanked by repetitive elements, and the longer-range information improves the mappability of the data, which provides more confident detection^36^. As the 3rd generation technologies mature, these structural variations could prove to be extremely significant to biomedicine and other analysis, as initial 3rd generation-based studies^8^ and older studies of copy number variations^37^ have suggested tens of thousands of structural variations, representing many millions of bases of sequence, are variable in a typical human genome compared to the standard reference, much of which is missed by short read sequencing.

**Haplotype phasing:** A fourth application is phasing heterozygous variants into separate haplotype-resolved sequences. This is important for analyzing allele-specific expression, determining parent of origin for *de novo* mutations, and other applications^38^. The phasing algorithms analyze the heterozygous variants in the genome, and use the read sequences or mapping information to link together the alleles that are present on the same chromosome. The analysis is complicated by sequencing errors and uneven coverages that can cause additional false variants to be introduced or true heterozygous variants to be missed. Consequently most algorithms use an optimization framework to improve robustness that assigns alleles to at most 2 possible haplotypes while minimizing the number of disagreements between the assignment and the underlying read information. In the case of the human genome, heterozygous variants occur on average every 1000bp to 1500bp making it unlikely that a short read will span two or more variants. In contrast, using ~5kbp Moleculo reads or ~100kbp 10X Genomics data, very large stretches of the genome can be robustly phased, and the published reports document haplotype blocks averaging more than 1Mb in length^19^.

The long-range information provided by the new sequencing and mapping technologies has primarily been applied to four main applications related to genome structure **(Figure 1 & Box)**. By carrying out a meta-analysis of the available 3^rd^ generation assemblies, and a large-scale simulation analysis of genomes across the tree of life we show how these technologies improve genome quality: near perfect assemblies are now possible for most organisms below 100Mbp in size, and very high quality assemblies are possible for larger genomes, including human and other mammalian genomes. We further compute a retrospective analysis of the improvements to the human reference genome, and show how the improved contiguity of the different builds has lead to an improved resolution of genomic features, such as to improve the number of gene clusters and the number of clinically significant variants that can be correctly identified. While such databases are currently only available in the well-studied human genome, we expect these types of results will become possible in other genomes with improved genome assemblies.

**Figure 1.**
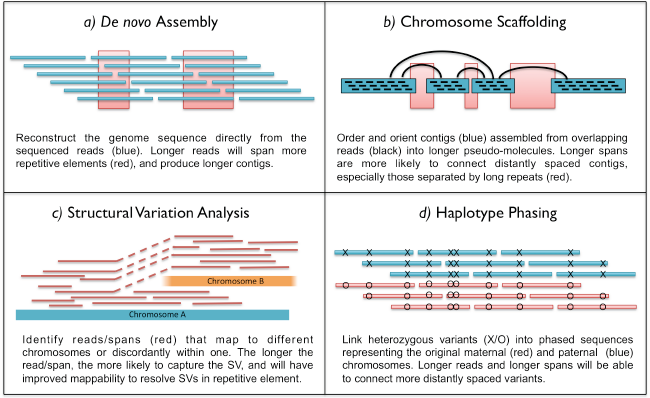
Overview of four major genomics applications empowered by long read/long span technologies.

### Contiguity

Contiguity is important to a genome assembly project so that genomic elements (exons, genes, gene clusters, transposons) are fully assembled, together with more contexts around each element, such as reconstructing complete genes along with their associated regulatory sequences. The Lander-Waterman statistics^39^, published in 1988, was one of the first and most widely used models of genome contiguity. For more than 25 years, it has guided researchers with useful recommendations of the minimum coverage needed for a sequencing project. It also predicts the average contig size that will be assembled from a given amount of coverage using reads of a certain length. However, the predictions become very poor at higher coverage, including predicting that the human genome should assemble into contigs hundreds of gigabases long with 100x coverage of 100bp reads, far beyond the length of the genome itself **(Online methods Figure 1)**.

The lack of predictive power in the Lander-Waterman statistics arises because it assumes that a genome is free of repetitive sequences, so that the reads can always be unambiguously assembled together. However, when applied to real genomes repeats cause ambiguity in how reads overlap, and genome assemblers will end contigs at repeats that are not spanned by sufficiently long reads^2^. Interestingly, a relatively modest increase in read length can exert a significant improvement to the assembly quality because of the exponentially decreasing repeat distribution found in many eukaryotic genomes **(Supplementary Figure S28, S29)**. For example, in the rice (*O. sativa*) genome, there are 300 times fewer exact repeats that are at least 3,650bp long (a typical Tru-Seq read length) compared to those that are at least 100bp long (a typical Illumina read length). Consequently, using the longer read technology can dramatically improve the assembly by spanning proportionally more of the repeats.

To build a more realistic model that accounts for the complexities of real genomes, we adopted a data driven approach using Support Vector Regression (SVR) that examined the composition of 26 different genomes ranging from the 1.66 Mbp *M. jannaschii* genome to the 3.0 Gbp *H. sapiens* genome **(Online Methods, Table S1)**. The genomes were selected to be a diverse, representative sample of genomes across the tree of life, consisting of 5 bacteria, 1 archae, 3 fungi, 1 amoebazoa, 8 plant, 3 invertebrate and 5 vertebrate species. Whenever multiple genomes of similar size were available, we selected the genome with the highest quality to ensure the analysis best captures the true complexities present. Notably, we excluded the very largest currently available genomes, such as the 22 Gbp Norway spruce, since the contiguities of those assemblies are poor (contig N50 < 50kbp), and would have distorted the analysis of the repeats present^40^.

For each of these genomes, we simulated shotgun sequencing them with varying read lengths and coverage **(Online Methods).** The average read lengths ranged from 3,650bp (mean1), 7,500bp (mean2) and 15kbp (mean4) to simulate third-generation sequencing technologies. We further doubled the read lengths three times to 30kbp (mean8), 60kbp (mean16) and 120kbp (mean32) reads, to simulate third-generation mapping technologies. For each genome and each read length, we simulated 5x, 10x, 20x and 40x coverage, and then assembled those reads with the Celera Assembler, which is the only published assembler currently available supporting reads longer than 100kbp. The reads were simulated without errors, representing the upper bound to the possible error correction strategies, although realistic parameters were used so that near identical repeats (within 2% similarity) will not be separated unless spanned by long reads, as would be used with genuine data^16^.

The results of the simulated read assemblies of the human genome as well as the N50 sizes of the best available published results using the different technologies are summarized in **Figure 2**. This figure, an N-chart, generalizes the N50 size and shows the contig/scaffold lengths sorted from longest to shortest. The red curve at the top shows the size distributions of the actual chromosome segments and bounds the possible results. For context, we also include the curves representing the scaffold and contig sizes from a genuine *de novo* assembly of the human genome using Illumina-only sequencing with ALLPATHS-LG^41^. For the third-generation technologies, we see strong overall agreement between the simulated and published results, although the published results lag the simulated results by a few percent, presumably because of the biases and systematic difficulties explained above.

**Figure 2.**
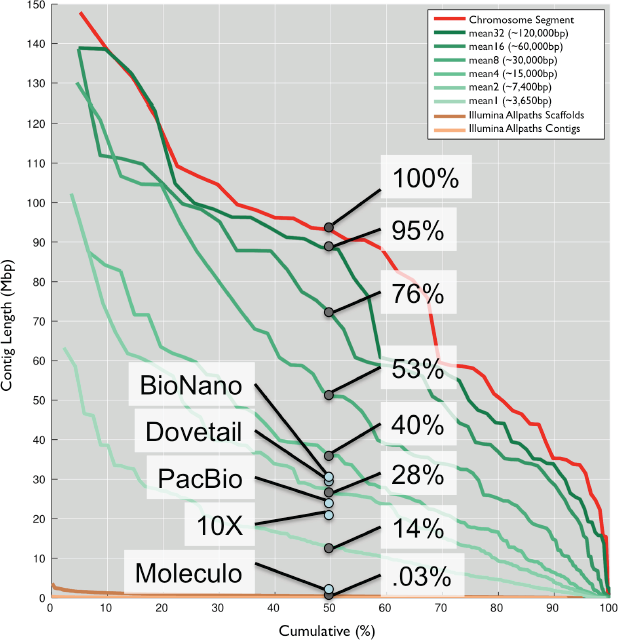
Contiguity of human genome assemblies. The red curve traces the lengths of the human chromosome segments and the green curves trace the results of different simulated read sets. The orange/brown curves trace the results of a de novo assembly of the human sample NA12878 using Illumina sequencing and ALLPATHS-LG (50x fragment coverage and 50x 2kbp mate pair coverage). The y-axis marks the length of the segment/contig, and the x-axis plots the cumulative fraction of the genome covered by segment/contigs that size or larger. The value at 50% marks the contig/scaffold N50 size. By construction the red curve has 100% assembly performance and the different simulated read sets have proportionally smaller percentages assembled. For context, the N50 size of several published human analyzes are also presented with blue circles as cited in Table 1.

The results across all 26 species, as well as a selection of available 3^rd^ generation assemblies shows assembly performance generally follows a logistic curve: the performance is consistently very high for small genomes, and drops off as the genome size increases depending on the read length used **(Figure 3)**. It is notable that with the current long read sequencing technologies (Illumina TruSeq, and error corrected PacBio and Oxford Nanopore sequences), the assembly performance is near 100% for most genomes less than 100Mbp in size, meaning it should be possible to assemble the complete chromosome arms of these species using the currently available technology. Indeed, very high quality and near perfect assemblies of several genomes of this size have been reported using these technologies. Beyond 100Mbp in size, the currently available read lengths should substantially improve assembly, and reach contig N50 sizes over 1 Mbp in many cases, although the achievable performance is still below entire chromosome segments unless long-range mapping technologies are applied.

**Figure 3.**
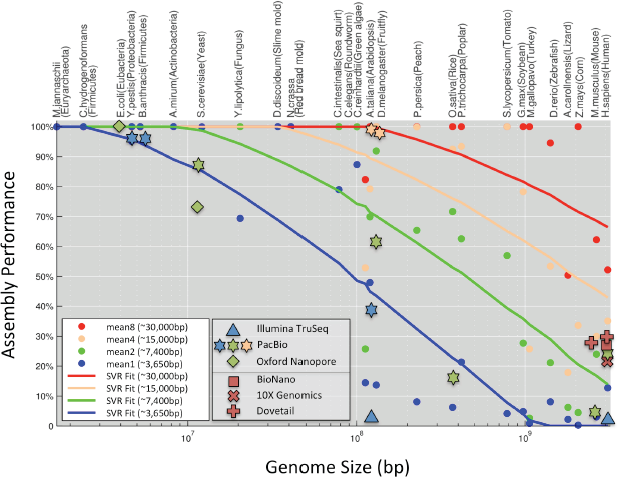
Assembly performance. The x-axis measures the genome size of the 26 genomes in log space. The y-axis measures the assembly performance of the different assemblies, meaning the N50 size of the assembly relative to the N50 size of the chromosome segments. Points indicate the results of simulated experiments with 20x coverage of error free reads of different read lengths. Lines show the best fit line from the SVR model from these simulated results. Other shapes indicate the genuine results of the assembly of real genomes using the different technologies, colored by their approximate equivalent simulated read lengths.

## Completeness

With sufficient sequencing coverage (>50x), every nucleotide of the genome should be sequenced, although the total span of the assembly will often nevertheless differ from that of the genome. This is because assemblers tend to filter out contigs shorter than a specified minimum length, mistakenly “collapse” repetitive sequences into fewer copies than are present in the genome, or introduce other artifacts^42^. Even the most recent builds of the human genome contain over one hundred million ‘N’s where repeats remain unresolved.

Gross assembly problems can be recognized if the total span of the assembly is substantially different from the true genome size, although assembly errors may both inflate and deflate the span of the assembled sequences by creating extra or fewer copies of sequences. A more focused gene-centric approach to assess completeness considers the fraction of genes or other genomic features that are completely and correctly assembled, using known genes, core eukaryotic genes, or *de novo* assembled transcripts for the evaluation.

To highlight the importance of a high quality assembly, we analyzed historical versions of the human genome, starting with the first published build from 2001, HG5, that had a 57kbp contig N50 size^5^ and ending with the more recent build, HG19 from 2009, that has a contig N50 size of 38Mbp (we denote contigs by the presence of Ns in the assembled sequence, not just chromosome segments). Although the different builds were not *de novo* assembled and used a combination of several new biotechnologies and extended read lengths, this analysis highlights how longer contigs translate into more complete assemblies. For each of the historical builds, we measured the fraction of genes annotated in HG19 that were fully intact in the older builds **(Figure 4, left)**. In addition to individual genes, we also considered blocks of 10, 100, or 1000 consecutive genes using a similar method to study overall genome organization. Only 93% of the individual genes of HG19 and less than 20% of 100 consecutive gene blocks were intact in HG5. It was not until the contig N50 size reached more than one megabase that almost all current genes and gene blocks were intact. We further evaluated the presence of known clinically relevant variations from ClinVar^43^ in the historical builds and find a substantial fraction (~10%) of these variations could not be recognized when the contig N50 size was less than 100kbp **(Figure 4, right)**. We expect this analysis to be a lower estimate on the significance of the improved contig sizes, especially as new technologies are used to discover more clinically relevant mutations in the currently “dark” regions of the genome^8, 36^.

**Figure 4.**
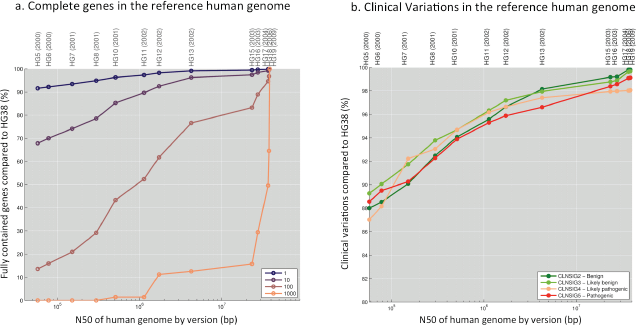
Completeness of historical human genomes. (left) Percentage of genes and gene blocks intact in historical build of the human genome. (right) Percentage of ClinVar clinically relevant variants present in the older builds of the human genome.

Finally, to study the relationship between completeness and contiguity across all species, we performed a similar gene block analysis for the different available assemblies of the 26 species **(Figure 5)**. Individual genes (gene blocks of length 1) were well captured in all assemblies while gene blocks of length 1000 were poorly captured the large genomes (>100Mbp). For gene blocks of intermediate lengths, such as blocks of 100 consecutive genes, the results follows logistic curve similar to the assembly performance contiguity curve. In this analysis, the enhanced genome completeness can be directly attributed to the improved read lengths used in the assemblies.

**Figure 5.**
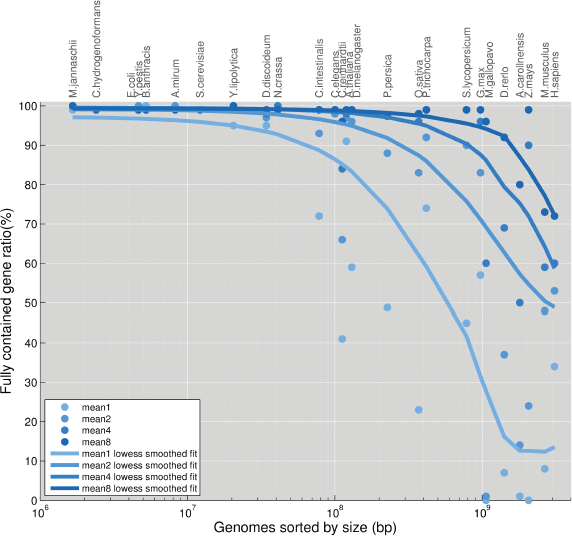
Gene Block Completeness of 26 genomes. For each of the 4 read lengths, we evaluated the fraction of 100 gene blocks annotated in each genome that were assembled completely intact. The solid lines represent the summary of the individual experiments computed with a local polynomial fit (lowess).

## Correctness

The correctness of a genome can be measured at the per-nucleotide or structural level. The per-nucleotide accuracy of assemblies using Illumina (or TruSeq) reads has been reported to be very high (> 99.9% accurate or higher) as it has very high nucleotide accuracy to start. Interestingly, despite their relatively high raw error rates (10% to 30% error), with sufficient coverage and proper algorithms both PacBio and Oxford Nanopore sequencing have produced assemblies with consensus nucleotide accuracy above 99.90%. For PacBio sequencing, the accuracy has been demonstrated to increase to 99.999% with increased coverage of just long PacBio reads^33^. This is because PacBio errors are dominated by random insertions and deletions, and it is increasingly unlikely that the same random mistake will occur at the same position in multiple reads^9^. To date, the accuracy of Oxford Nanopore assemblies, using either hybrid correction with short reads or pure-Oxford Nanopore assemblies, has lagged behind the PacBio results. This is largely because of the increased rate of non-random errors, especially in homopolymer sequences, although improvements to the sequencing technology and analysis algorithms currently in development are expected to improve the accuracy. Per-nucleotide accuracy is generally unaffected by long-range mapping since these technologies are primarily used to order and orient existing sequences, not to add or replace them.

In contrast to per-nucleotide accuracy, most common structural errors occur because of repetitive sequences in the genome, which can result in the “collapse” of a repeat into a single occurrence, or the reordering of sequences that occur between repeat copies^42^. These types of errors occur when the boundary of repeats is not recognized, especially at low coverage levels when the presence of a repeat may go undetected^44^. Longer reads are useful for reducing the frequency of structural errors as they span more repeats in the genome. For example, there is more than an order of magnitude decrease in major mis-assemblies present in the human genome assembled with synthetic 150kbp reads (mean32) versus 3,600bp reads (mean1) (**Figure 6**). Similar trends are observed in all the other genomes as the read lengths improve, although the smallest genomes are perfectly assembled by even the shortest reads considered.

**Figure 6.**
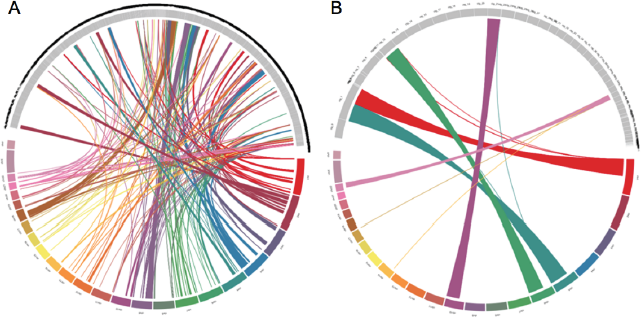
Human assembly structural correctness.

(left) The *de novo* assembly of with 20x coverage of the mean1 reads is shown at the top half of the circle and the reference human genome (hg19) is shown at the bottom. Colored bars show large-scale mis-assemblies where an assembled contig is mapped to two or more chromosomes. (right) The *de novo* assembly of 20x coverage of the mean 32 reads is displayed in a similar representation. For clarity, alignments of contigs that correctly align to a single chromosome are not displayed.

## Discussion

Third-generation DNA sequencing and mapping technologies are starting to produce genome sequences of remarkable quality. This is most easily measured by the contig and scaffold N50 sizes reported using these technologies that are hundreds to thousands of times more contiguous than corresponding short read assemblies. These assemblies, with megabase contigs and multi-megabase scaffolds, are truly reference quality and enable improved analysis of nearly every aspect of a genome: more complete and accurate representations of genes, clinically relevant SNPs, regulatory regions and other important genomic elements, as well as better resolution of the overall chromosome organization.

The highest quality genomes available have been assembled from the longest possible reads aided by the longest possible mapping information. Interestingly, the per-nucleotide error rate of the reads have had little effect on the per-nucleotide assembled sequence accuracy, as well-tuned algorithms can effectively reduce even 30% per-nucleotide error to below 1% with sufficient coverage. Our meta-analysis of available assemblies together with the modeling we present shows that it should be possible to assemble nearly complete chromosomes for genomes of up to 100Mbp in size with the currently available long-read technologies. For larger genomes, great gains are possible over strictly short read sequencing, with results approaching or exceeding those from older, more expensive BAC-by-BAC or fosmid-based assemblies. If the project demands even higher quality assemblies, the model also forecasts when those data may be available. In particular, for the human genome the read lengths need to average over 150kbp before complete chromosomes should be possible. If the historical trends continue, this could be achieved in as little as 3 to 4 years. When that milestone is reached, it is likely that many projects will begin from the fully assembled genomes instead of variant lists, opening new opportunities for studying structural variations across large populations^45^.

Our analysis includes assembling simulating reads from published reference genomes as an upperbound of their utility: the genomes we analyzed have gaps and errors that mask their true complexity, and our simulated reads do not contain errors nor any heterozygosity. In practice, researchers may need to oversample the genome more than predicted to account for any residual errors or biases present. Indeed, while our analysis suggests that 20x coverage of a genome should be enough to well assemble a genome, we recommend researchers sample >75x when using the new long read sequencing technologies to make the error correction steps most effective and to ensure high coverage is available of the longest reads. Ideally, if the budget and sample materials allows, we recommend assembling 20x coverage of error corrected reads exclusively over 20kbp long, using haploid or inbred samples if possible. We also caution researchers to carefully monitor the developments to the field as all of these technologies are rapidly evolving and new technologies are already under development. Both PacBio and Oxford Nanopore have announced higher throughput and lower cost instruments will be released this year, and the new 10X and Dovetail technologies are rapidly improving.

## Abbreviations

BAC: Bacterial Artificial Chromosome; bp: base pair; kbp: kilobases; Gbp: gigabases; Mbp: megabases; SNP: single nucleotide polymorphism.

## Acknowledgements

This project was supported in part by National Science Foundation awards DBI-1265383 and DBI-1350041 to MCS, and IOS-1032105 and DBI-0922738 to WRM. It was also supported in part by National Institutes of Health award R01-HG006677 to MCS. Also this work was done in part while HL was visiting the Simons Institute for the Theory of Computing, University of California, Berkeley We would like to thank Adam Phillippy, Sergey Koren, Jason Chin, Paul Peluso, David Rank, Wendy Pepper Weise, and Greg Khitrov for their helpful discussions.

## Contributions

H.L performed computational experiments on assembly simulation, data analysis and modeling, and wrote the manuscript. J.G and M.N. performed the non-simulated assemblies and wrote the manuscript. S.Y contributed to modeling and prediction using machine learning and edited manuscript. S.M and S.G. performed data analysis and wrote the manuscript. W.R.M and M.C.S designed the study, supervised the project, and wrote the manuscript.

